## Supplementary material for "Analysis of computational codon usage models and their association with translationally slow codons": S1 GWIPS-viz Data Information

The following table contains information about the 17 RFP data sets we analyzed from GWIPS-vis. The first column (“Track”) relays the original study, the second (“Table”) indicates the subtrack (or specific data set) used, as many studies have multiple published data sets associated with them, and the third column indicates whether or not that data set has replicates. Track and table names are taken directly from GWIPS-vis. These data sets can be downloaded at gwips.ucc.ie by first clicking “Downloads”, then setting the Clade to “Yeast”, the Genome to “S. cerevisiae”, the Assembly to “Apr. 2011 (SacCer_Apr2011/sacCer3”, the Group to “Elongating Ribosomes (A-site)”, then selecting the corresponding Tracks and Tables listed below.

| Track | Table | Replicates? |
| --- | --- | --- |
| Albert 2014 | BY FP | No |
| Cai 2013 | Ribosome footprints for wildtype | No |
| Guydosh 14 | wild_type_no_additive | No |
| Guydosh 14 | wild_type_CHX | No |
| Ingolia 2009 | Rich | Yes |
| Lareau 2014 | Read length 25+: Untreated | Yes |
| Lareau 2014 | Read length 25+: Cycloheximide | Yes |
| Nedialkova15 | WT ribo YPD | Yes |
| Nedialkova15 | WT ribo YPD noCHX | Yes |
| Nissley 2016 | yeast S288C Riboseq | Yes |
| Pop 2014 | WT footprint | No |
| Schmidt16 | Control RPF | No |
| Subtelny 2014 | Cerevisiae RPF | No |
| Thiaville 2016 | RiboSeq WT | Yes |
| Young 15 | wild_type | No |
| Zid 2014 | BYRibo_logphase | Yes |
| Zinshteyn 2013 | WT Ribosome Footprint | Yes |

For further information about each of the data sets and the studies that originated them, click on “Genome Browser” from the GWIPS-vis home page, then make sure the selected species is “S. cerevisiae” and the assembly is “Apr. 2011 (SacCer_Apr2011/sacCer3”, then click “Go”. Scrolling down on the page will find a link for every study that has data hosted on GWIPS-vis (note that we use the Group “Elongating Ribosomes (A-site)”. By clicking on any of the studies, you can find out more information about it, as well as a reference to the original paper at the bottom of the page.
